# DNA entry into the cohesin ring through the kleisin N-gate

**DOI:** 10.64898/2026.08.04.742744

**Authors:** Karolina Gmurczyk, Torahiko L. Higashi, Georgia Whitton, Frank Uhlmann

## Abstract

The ring-shaped cohesin complex topologically entraps two DNAs to establish sister chromatid cohesion, a prerequisite for faithful chromosome segregation during cell divisions. How DNA enters the cohesin ring remains incompletely understood. Both the interface between Psm3^Smc3^ and the Rad21^Scc1^ N-terminus - the N-gate - and the SMC hinge interface - the hinge gate - can open *in vitro* to entrap DNA. Here, we combine biochemical with *in vivo* functional assays in the fission yeast *S. pombe* to probe the DNA entry mechanism. Locking the hinge gate does not interfere with cohesin loader-and ATP hydrolysis-dependent DNA entrapment. Conversely, cohesin with a tightly locked N-gate entraps DNA *in vitro* but does not load onto chromatin or establish sister chromatid cohesion *in vivo*. The cohesin ATPase is functionally coupled to N-gate, but not hinge-gate, opening. Our results put forward the N-gate as the primary DNA entry gate into the cohesin ring.

## INTRODUCTION

Two sister chromatids must be held together from their replication in S phase to their segregation in anaphase. The spatial proximity of sister chromatids, called sister chromatid cohesion, is crucial to maintaining genomic stability through cell divisions. Sister chromatid cohesion is established by a multisubunit protein complex, cohesin. Cohesin is a conserved, ring-shaped assembly made up of the Psm1, Psm3, Rad21 and Psc3 subunits in fission yeast (known as Smc1, Smc3, Scc1 and Scc3 in budding yeast and SMC1, SMC3, RAD21, and STAG1/2 in humans) ^1–4^. The two SMC subunits Psm1 and Psm3 together with the kleisin subunit Rad21 form the ring circumference, while Psc3 interacts with the ring through Rad21 (Figure 1A). This ring architecture allows cohesin to entrap DNA by topological embrace ^5–9^.

**Figure 1.**
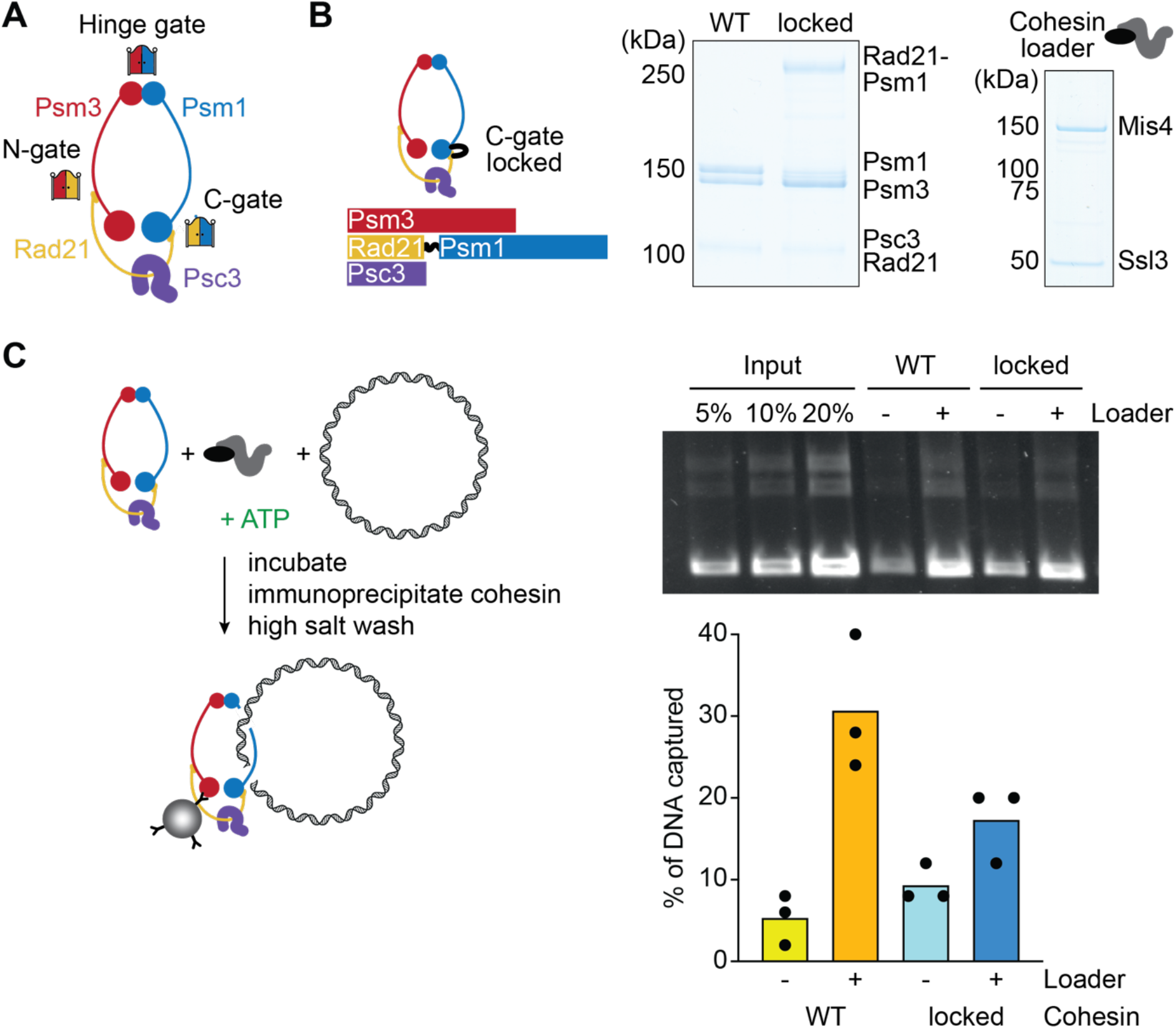
*In vitro* DNA entry into C-gate locked cohesin. (A) Schematic of the cohesin tetramer complex and its three potential DNA entry gates. (B) Schematic of a C-gate locked cohesin complex, as well as purified WT and C-gate locked cohesin complexes analysed by SDS-PAGE and Coomassie Blue staining. The purified Mis4-Ssl3 cohesin loader complex was analysed alongside. (C) Experimental diagram of the cohesin loading assay and representative agarose gel image of DNA recovered in a cohesin loading assay using WT and C-gate locked cohesin, with and without added cohesin loader. Recovered DNA was quantified in three independent experiments. The individual results are shown, bars represent the means.

Cohesin complexes assemble before loading onto DNA ^10^. Therefore, at least one of the three protein-protein interfaces between cohesin’s ring subunits, known as its ‘gates’, need to transiently open to allow DNA entry into the cohesin ring. The ‘N-gate’ forms between the Rad21 N-terminus and the Psm3 neck, the coiled coil region adjacent to the Psm3 ATPase head; the ‘C-gate’ forms between the Rad21 C-terminus and the Psm1 ATPase head; while the ‘hinge gate’ lies between the Psm1 and Psm3 halves of the hinge domain (Figure 1A) ^11–16^. Past analyses have led to the suggestion that the C-gate is not involved in topological cohesin loading onto DNA ^17,18^. What remains less clear is whether the N-gate or the hinge gate serves as the primary DNA entry gate. *In vitro*, cohesin can entrap DNA by using either the N-gate or the hinge gate ^19^. DNA entry through the N-gate finds support from a detailed structure-based mechanism, in which the N-gate is first opened upon ATP binding and then closed as the cohesin loader, Mis4, locks the DNA against the ATPase heads. Next, ATP hydrolysis disengages the ATPase heads to allow completion of DNA entry ^15,20,21^. Consistent with DNA entry through the N-gate, *Xenopus* cohesin with a chemically, covalently locked hinge gate remains proficient in loading onto chromatin ^22^. On the other hand, evidence has also been presented that DNA apparently enters the cohesin ring through the hinge gate. Hinge gate closure, using additionally inserted protein dimerization domains, blocks cohesin loading onto chromatin in yeast and human cells. In turn, fusing the Smc3 C-terminus to the kleisin N-terminus by a flexible linker, an approach interpreted as locking the N-gate, did not block cohesin loading onto DNA^17,23,24^.

Meanwhile, cryo-EM structures of cohesin have revealed the molecular context of cohesin’s gates, and they have provided a better understanding of the methods that were designed to lock them ^15,16,18,20^. The linkers used to fuse the Smc3 C-terminus to the kleisin N-terminus, intended to lock the N-gate, were disproportionately long ^17,24,25^. Such long linkers should not in fact interfere with the operation of the N-gate, nor would they prevent DNA entry through this gate. Rather, DNA that enters through the ‘locked’ N-gate would be accommodated in an additional compartment that is formed by the linker peptide chain ^26^. With regards to the hinge gate, two structural studies revealed that the hinge intimately engages with both Psc3 and the cohesin loader Mis4 ^15,16^. These interactions make the hinge fold back, in turn making the N-gate accessible to oncoming DNA. Dimerization domain insertions at the hinge, especially once locked ^17,19,23^, might have compromised these Psc3 and Mis4 interactions. Thus, the hinge insertions could have inadvertently, but indirectly, affected DNA entry through the N-gate.

Based on the above considerations, we here revisit the cohesin gate closure experiments. We design updated versions of C-gate, N-gate, and hinge gate locked cohesin, and we test their ability to entrap DNA *in vitro*, as well as their abilities to load onto chromatin and establish sister chromatid cohesion *in vivo*. We find that cohesin with a covalently locked hinge gate entraps DNA *in vitro* in an ATP-and cohesin loader-dependent manner indistinguishable from wild type cohesin. In comparison, tightly locking the N-gate reduces *in vitro* DNA entrapment and abolishes cohesin’s ability to load onto chromatin and perform sister chromatid cohesion. We also find that N-gate, but not hinge gate, opening is coupled to cohesin’s ATPase. Our results put forward the N-gate as cohesin’s primary DNA entry gate.

## RESULTS

### *In vitro* DNA entry into cohesin with a locked C-gate

Even when its C-gate is closed by a covalent Scc1-Smc1 peptide linker, budding yeast cohesin topologically entraps DNA *in vitro*, and loads onto chromatin *in vivo* ^17,19,22^. To investigate whether this behaviour is conserved amongst species, we turned to the fission yeast *S. pombe*. We purified fission yeast cohesin tetramers (consisting of Psm3, Psm1, Rad21 and Psc3), as well as a variant in which the C-gate was locked by fusing Rad21 to Psm1 by a peptide linker (Figures 1B and S1A). We also purified the Mis4-Ssl3 cohesin loader dimer that stimulates topological cohesin loading onto DNA ^1,8^. To test the ability of WT and C-gate locked cohesin to topologically entrap DNA, we incubated these cohesins with circular DNA, ATP, and with or without the cohesin loader. Cohesin was immunoprecipitated using magnetic beads, washed with high salt buffer to disrupt non-topological cohesin-DNA interactions, then the recovered DNA was visualised by agarose gel electrophoresis (Figure 1C). This setup reports on topological DNA entrapment, as no DNA was recovered following incubation with linear DNA (Figure S1B). Similarly to wild type cohesin, C-gate locked cohesin entrapped DNA poorly in the absence of the cohesin loader. Cohesin loader addition increased DNA recovery by both WT and C-gate locked cohesins, though C-gate locked cohesin entrapped somewhat lower DNA levels compared to wild type cohesin. We note that C-gate locked budding yeast cohesin also showed reduced *in vitro* DNA entrapment in two complementary topological DNA entrapment assays ^19,22^. These results show that DNA entry into the cohesin ring is possible *in vitro* when the C-gate is locked, albeit at a reduced efficiency.

### *In vivo* chromatin loading and sister chromatid cohesion establishment by C-gate locked cohesin

We addressed if C-gate locked fission yeast cohesin can load onto chromatin and establish sister chromatid cohesion *in vivo*. To do so, we used a *rad21-k1* temperature sensitive strain background ^27^, into which we inserted ectopic genes encoding either wild type Rad21, or a C-gate locked Rad21-Psm1 fusion protein. (Figure 2A). The Rad21-Psm1 fusion protein rescued the viability of the *rad21-k1* strain at its restrictive temperature to a large degree (Figure 2B), suggesting that it formed part of functional cohesin complexes.

**Figure 2.**
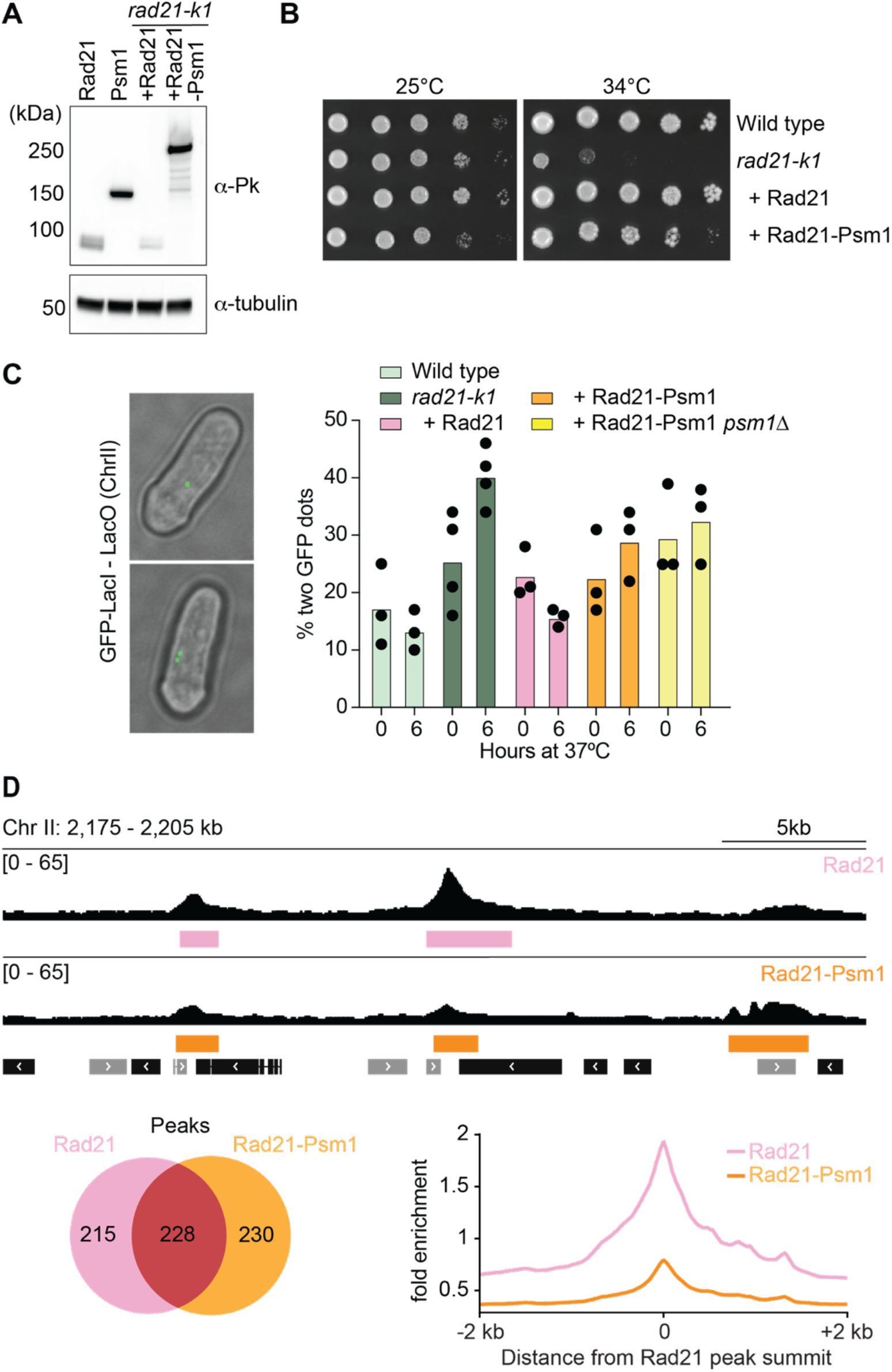
In vivo chromatin loading and sister chromatid cohesion establishment by C-gate locked cohesin. (A) Ectopic Rad21 and Rad21-Psm1 expression levels, genes inserted at the *arg1^+^*locus in the *rad21-k1* strain background (right lanes), next to endogenous Rad21 and Psm1 levels (left lanes), all tagged with Pk_3_ epitope tags and analysed by Western blotting. Tubulin served as a loading control. (B) Cells of the indicated genotypes were spotted at 10-fold serial dilutions, and cell viability was assessed at 25°C and 34°C, permissive and restrictive temperatures for the *rad21-k1* allele,respectively. (C) Example images of cells showing cohesed or separated sister chromatids, visualised by LacI-GFP bound to LacO sequences inserted close to the centromere of chromosome II. Quantification of sister chromatid cohesion is shown alongside in three biological repeat experiments of the indicated strains, before and 6 hours after shift to 37°C. The individual results are shown, bars represent the means. (D) Calibrated ChIP sequencing traces of Rad21 and Rad21-Psm1 along a representative region of chromosome II, with detected peaks demarcated. The positions and orientation of protein coding genes are indicated. The Venn diagram represents detected peak numbers and their overlap, while a peak average plot was generated around the centres of the 228 shared peaks.

To examine the proficiency of C-gate locked cohesin to establish sister chromatid cohesion, we used strains in which a locus close to the centromere of chromosome II was marked with a lac operator array, visualised by a lac repressor-GFP fusion protein ^28,29^. Two sister chromatids that are held together by cohesin are visible as a single GFP dot, while defective sister chromatid cohesion results in two separated GFP signals (Figure 2C). We shifted cells to 37°C for six hours before assessing sister chromatid cohesion. Fission yeast cells spend the majority of their growth and division cycle in G2 ^30^, and a fraction of cells is in the process of chromosome segregation, resulting in a background level of two GFP dots even in wild type cells. Above that background, *rad21-k1* cells showed a pronounced sister chromatid cohesion defect. This defect was rescued by expression of wild type Rad21, and partially by the C-gate locked Rad21-Psm1 fusion protein.

Cells expressing the Rad21-Psm1 fusion protein additionally harboured endogenous wild type Psm1. Therefore, cohesin complexes could form that contain Rad21 from the Rad21-Psm1 fusion, bound to wild type Psm1. To eliminate this possibility, we deleted the wild type *psm1^+^* gene. Following *rad21-k1* inactivation, the Rad21-Psm1 fusion protein now was the only source of Rad21 and Psm1. This deletion did not result in any further sister chromatid cohesion defects (Figure 2C), confirming that C-gate locked cohesin sustained cell growth and sister chromatid cohesion.

Next, we examined chromatin association of C-gate locked cohesin using calibrated chromatin immunoprecipitation and sequencing (ChIP-seq). We cultured *rad21-k1* cells expressing Rad21 or Rad21-Psm1 at 25°C and immunoprecipitated the ectopically expressed proteins by virtue of their Pk epitope tags. Three biological repeat experiments showed good reproducibility (Figure S2A) and were therefore aggregated. We detected 443 Rad21 peaks and 461 Rad21-Psm1 peaks, 228 of which overlapped (Figure 2D). Most peaks uniquely enriched for Rad21 were at sites of convergent transcription, as expected ^31–34^, while most peaks uniquely enriched for Rad21-Psm1 colocalised with binding sites of the Mis4-Ssl3 cohesin loader and promoters ^35^. Peaks common to both Rad21 and Rad21-Psm1 were distributed between cohesin loader binding sites, promoters and sites of convergent transcription (Figure S2B). Aggregate peak profiles of the 228 peaks shared between Rad21 and Rad21-Psm1 revealed similar peak characteristics, yet lower Rad21-Psm1 levels compared to wild type Rad21 (Figure 2D). Taken together, our results demonstrate that C-gate locked cohesin loads onto chromatin at somewhat reduced levels, with some alterations to its chromatin distribution that we will discuss further below. Overall, locking the C-gate remains compatible with cohesin’s roles and functions in fission yeast.

### *In vitro* DNA entry into cohesin with a locked hinge gate

We next turned our attention to the hinge gate, given conflicting results as to whether it serves as a DNA entry gate ^15,17,19,22^. We purified the cohesin tetramer containing CLIP-and SNAP-tag insertions in the Psm3 and Psm1 halves of the hinge, respectively (Figure 3A). These two insertions can be covalently linked by a bifunctional crosslinker in which a SNAP substrate is linked to a CLIP substrate via a Cy5 dye moiety (SC-Cy5) ^15,36^. After the crosslinking incubation, approximately half of cohesin complexes had a covalently closed hinge gate, as seen during SDS polyacrylamide gel electrophoresis (Figures 3A and S3A). In our topological DNA loading assay, the crosslinked cohesin entrapped DNA with equal efficiency as cohesin that was left untreated (Figure 3B), suggesting that closing the hinge gate had not impeded DNA entry into the cohesin ring.

**Figure 3.**
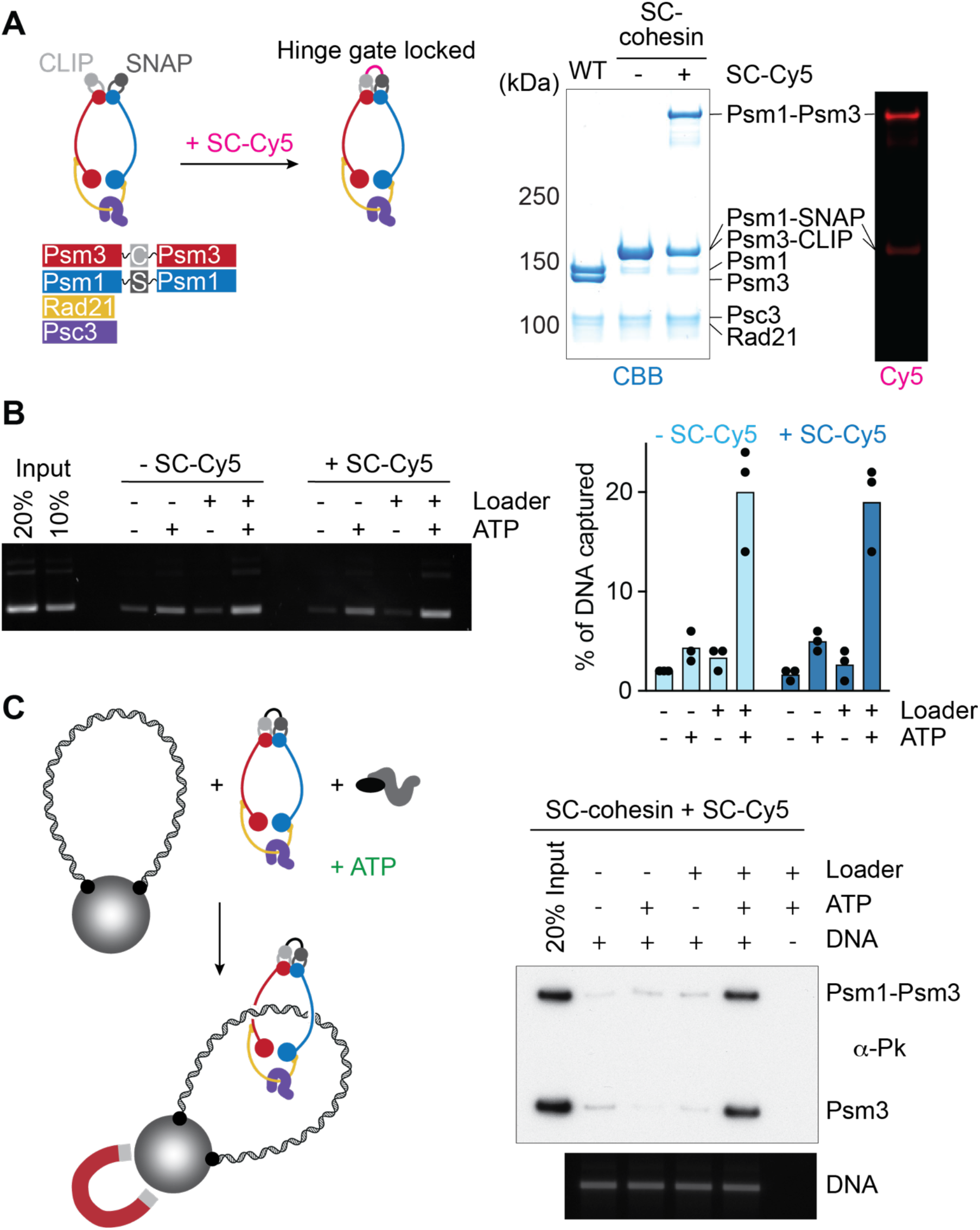
In vitro DNA entry into hinge gate locked cohesin. (A) Schematic of a hinge gate locked cohesin complex, using the bivalent SC-Cy5 crosslinker that covalently bridges the SNAP and CLIP tag insertions. Purified WT, as well as untreated and SC-Cy5 treated hinge gate locked cohesin complexes were analysed by SDS-PAGE and Coomassie Blue staining. Cy5 in gel fluorescence is shown alongside. (B) Representative agarose gel image of DNA recovered in a cohesin loading assay using untreated or SC-Cy5 treated hinge gate locked cohesin, as well as quantification of 3 independent repeat experiments. The individual results are shown, bars represent the means. (C) Schematic of cohesin loading onto topologically closed DNA, immobilised on magnetic beads. A representative Western blot image of loaded cohesin, with its hinge gate open or closed, and an agarose gel image of the recovered DNA are shown.

A confounding factor to the conclusion that hinge crosslinking did not impede DNA entry is the fact that SC-Cy5 covalently closed the hinge gate of only half of the cohesin complexes. We therefore used an alternative approach to investigate whether, and how, cohesin with a covalently closed hinge gate entraps DNA. To do so, we tethered two ends of linear DNA to magnetic beads to create an immobilised, topologically closed DNA substrate ^15^. We incubated this beads-DNA substrate with SC-Cy5 treated cohesin, ATP and the cohesin loader. After pull-down and high salt washes we divided the reaction into two, to visualise both the bead-bound DNA and cohesin that had loaded onto it (Figure 3C). This setup revealed that both cohesins, with or without a covalently closed hinge, entrapped DNA equally efficiently. DNA entrapment was topological in nature, as restriction enzyme cleavage resulted in cohesin release from the DNA (Figure S3B). Therefore, DNA does not need to pass through the hinge gate to enter the fission yeast cohesin ring.

We furthermore investigated the ATP and cohesin loader requirements for the loading of hinge-gate locked cohesin onto DNA. ATP and the cohesin loader stimulated hinge-gate locked and wild type cohesin loading onto DNA in equal measures (Figure 3C). These results suggest that the reaction by which cohesin loads onto DNA when the hinge gate is locked, i.e. most likely through the N-gate, shows the same requirements for ATP and the cohesin loader as does cohesin loading onto DNA *in vivo* ^1,37,38^.

Experiments with budding yeast cohesin also found that hinge gate locked cohesin rings topologically load onto DNA. However, in that case, the reaction was not stimulated by the cohesin loader ^19^. The difference to our results could reflect inter-species differences, or they might be explained by the different dimerization tags used (the SpyTag-SpyCatcher system ^39^), or the places where they were inserted. The hinge folds back by engaging in interactions with both Psc3 and the cohesin loader, to facilitate DNA access to the N-gate ^15,16,40^. It is possible that certain hinge insertions interfered with these interactions and thereby, while the insertions were made at the hinge, in fact compromised DNA entry through the N-gate (Figure S3C).

### *In vivo* locking of the hinge gate

Having established that locking the hinge gate is inconsequential for *in vitro* DNA entry into the fission yeast cohesin ring, we wanted to study the possible consequences of hinge gate closure on *in vivo* cohesin function. We utilised the third generation SpyTag-SpyCatcher pair, reported to efficiently create a spontaneous isopeptide bond between its two components ^39,41^. Individual insertions of the Spytag into Psm3 (Psm3-ST) and of the SpyCatcher at two places into Psm1 (Psm1-SC) were tolerated well, but their combination as the sole source of these SMC subunits did not support cell growth or the establishment of sister chromatid cohesion (Figure S4). However, the inability of Psm3-ST and Psm1-SC to provide cohesin function was unlikely due to hinge gate closure, as protein size analysis revealed that isopeptide bonds formed only in a small fraction of cohesin complexes. We ascertained that the small fraction of hinge gate closure did not exert a dominant negative effect, as closed cohesin was well tolerated as long as wild type Psm1 was also present. Thus, we were unsuccessful with efficiently locking the hinge gate and studying the consequences *in vivo*. Instead, our efforts underscore the sensitivity of the hinge, the cohesin ring’s weakest subunit interface ^42^, to perturbations.

### *In vitro* DNA entry into cohesin with a locked N-gate

To explore if fission yeast cohesin with a locked N-gate can entrap DNA, we purified two types of cohesin complexes containing Psm3-Rad21 fusions (Figure 4A and S5A). The first type included a 60 amino acids long linker between Psm3 and Rad21 (Psm3-L-Rad21), similar in length to previously studied budding yeast and human N-gate locked cohesin complexes ^17,23^. Generating a Psm3-L-Rad21 structural model shows that the linker length exceeds what would be necessary to connect Psm3 to Rad21 (Figure 4B). We therefore generated a Psm3-S-Rad21 fusion protein with a shorter, 14 amino acids linker. This linker length should readily connect the structured components of both subunits, without adding unnecessary slack.

**Figure 4.**
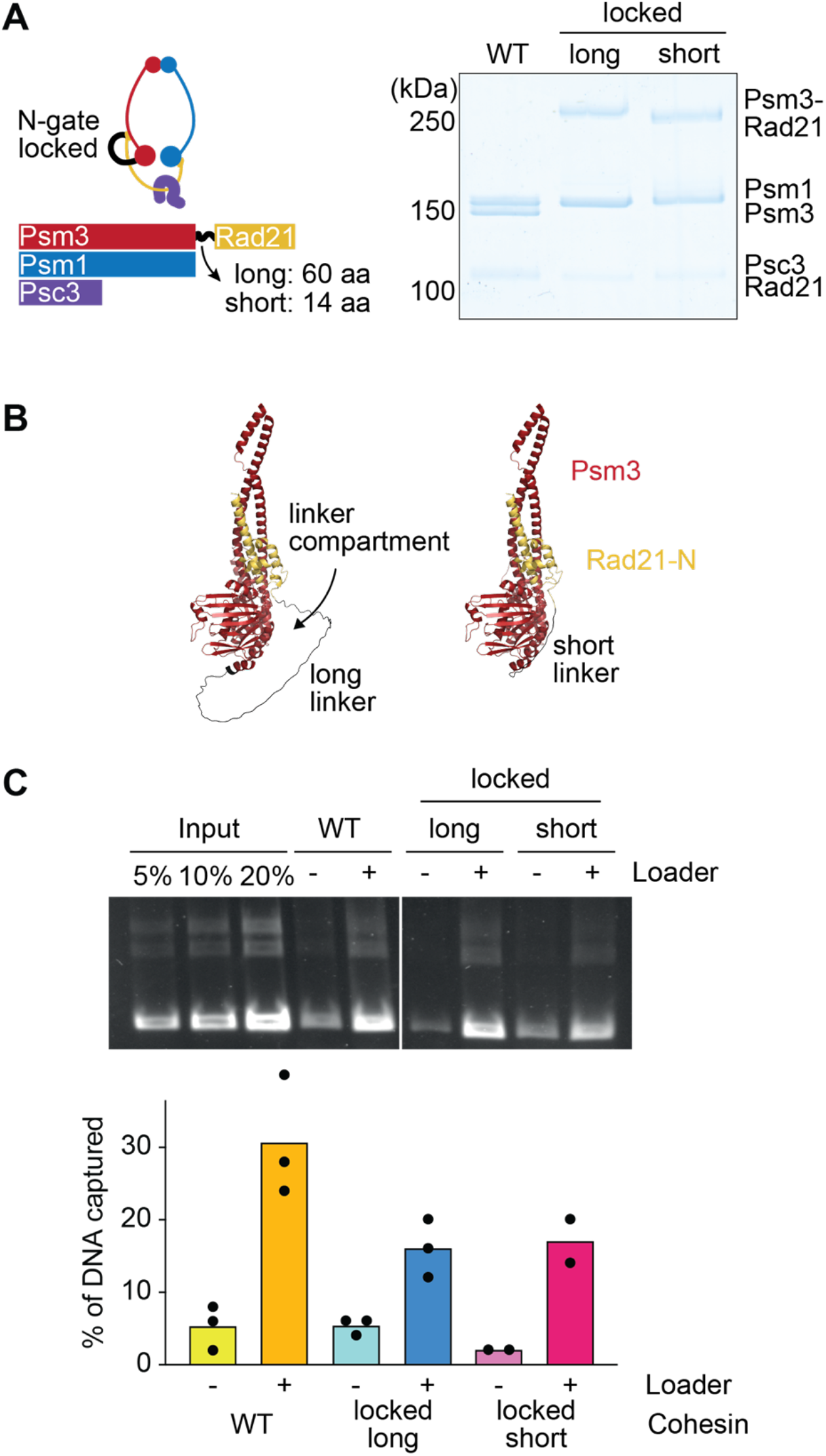
In vitro DNA entry into N-gate locked cohesin. (A) Schematic of an N-gate locked cohesin complex, with the long and short linker lengths indicated, as well as purified WT and the two N-gate locked cohesin variants analysed by SDS-PAGE and Coomassie Blue staining. (B) Structural models of the Psm3-L-Rad21 (long) and Psm3-S-Rad21 (short) fusions. The linker compartment generated by the long linker is indicated. Models were generated using AlphaFold 3 ^61^. (C) Representative agarose gel image of DNA recovered in a cohesin loading assay comparing WT and the two N-gate locked cohesin variants, as well as quantification of 2 or 3 independent repeat experiments. The individual results are shown, bars represent the means.

We next tested the ability of both types of N-gate locked cohesin to entrap DNA *in vitro*. Psm3-L-Rad21 and Psm3-S-Rad21 cohesin could both be loaded onto DNA, in reactions that were stimulated by the cohesin loader, albeit at reduced efficiencies when compared to wild type cohesin (Figures 4C). Therefore, a covalent linkage between Psm3 and Rad21 does not prevent DNA entry into the cohesin ring. These results agree with data obtained with *S. cerevisiae* cohesin, where N-gate closure did not block DNA entry into the cohesin ring ^19^. We note that our assay probed the topological entrapment of DNA by the cohesin ring, but did not address where in the cohesin ring the DNA resides following loading.

### A tightly locked N-gate stops cohesin function

To test whether a locked N-gate is compatible with *in vivo* cohesin function, we expressed Psm3-L-Rad21 and Psm3-S-Rad21 in a temperature sensitive *psm3-602* fission yeast strain background, at levels comparable to endogenous Psm3 and Rad21 (Figure 5A). Strikingly, Psm3-L-Rad21, but not Psm3-S-Rad21, supported cell growth at 34°C, a restrictive temperature of the *psm3-602* allele (Figure 5B). Furthermore, Psm3-L-Rad21 partially restored the sister chromatid cohesion defect seen in the *psm3-602* strain, but Psm3-S-Rad21 did not (Figure 5C). These results show that fusing Psm3 with Rad21 using a long linker creates a functional cohesin complex, consistent with what was seen in budding yeast ^17^. Reducing the linker length, while not adversely affecting cohesin’s *in vitro* measurable activities, stopped cohesin’s ability to support cell viability and sister chromatid cohesion.

**Figure 5.**
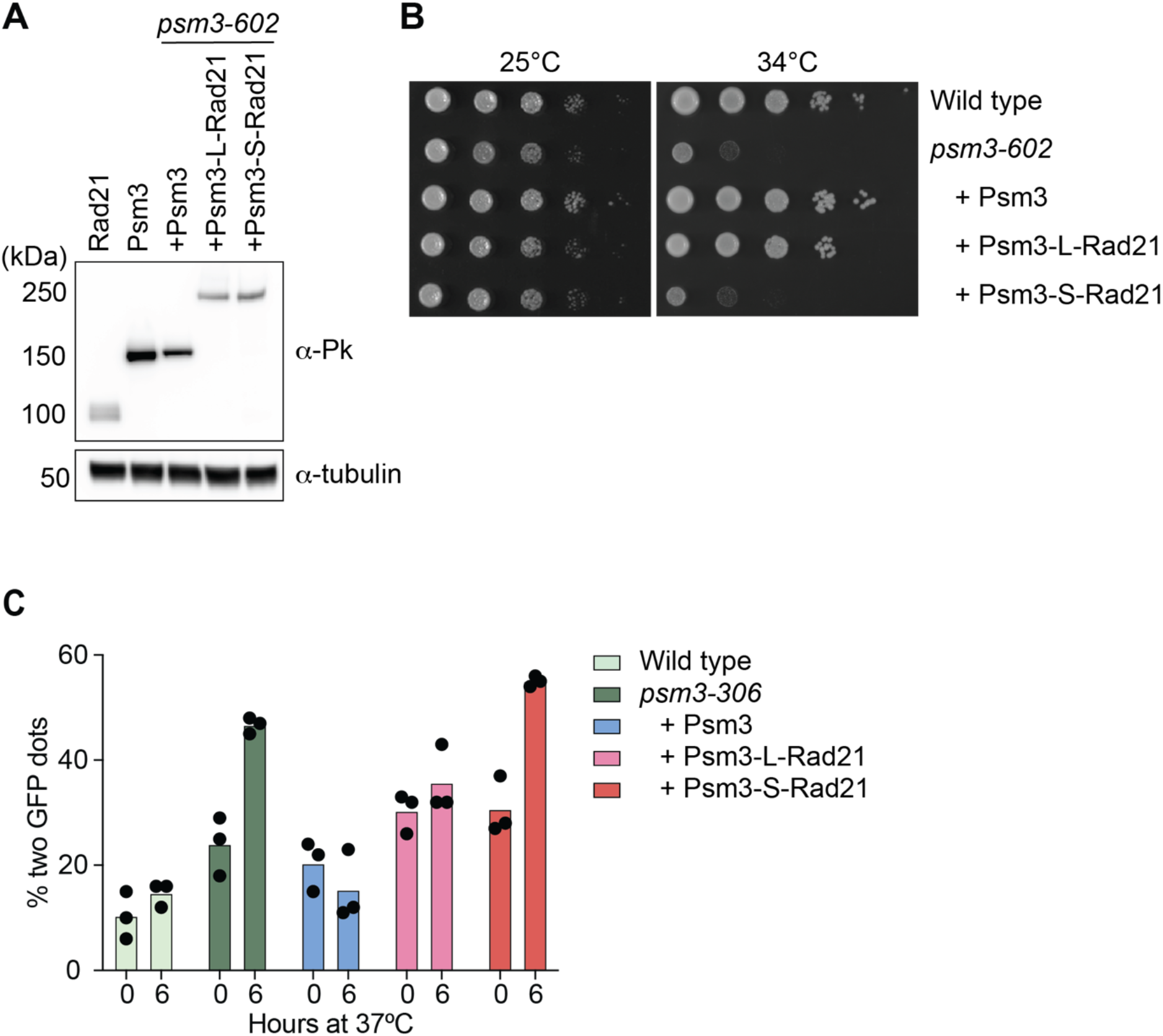
In vivo sister chromatid cohesion establishment by N-gate locked cohesin. (A) Ectopic Psm3, Psm3-L-Rad21, and Psm3-S-Rad21 expression levels in the *psm3-602* strain background (right lanes), next to endogenous Rad21 and Psm3 levels (left lanes), all tagged with Pk_3_ epitope tags and analysed by Western blotting. Tubulin served as a loading control. (B) Cells of the indicated genotypes were spotted at 10-fold serial dilutions, and cell viability was assessed at 25°C and 34°C, permissive and restrictive temperatures for the *psm3-602* allele, respectively. (C) Quantification of sister chromatid cohesion defects, seen as two separated GFP dots, in three biological repeat experiments of the indicated strains, before and 6 hours after shift to 37°C. The individual results are shown, bars represent the means.

### A tightly locked N-gate prevents *in vivo* cohesin loading

To investigate the ability of the two N-gate locked cohesin variants to load onto chromatin, we turned to our quantitative ChIP sequencing approach. We cultured *psm3-602* cells expressing WT Psm3, Psm3-L-Rad21, or Psm3-S-Rad21, each fused to the same Pk epitope tag to facilitate pulldown. At least three biological replicates of the ChIP experiments were performed and aggregated (Figure S5). We identified 497 Psm3 peaks and 644 Psm3-L-Rad21 peaks, around half of the latter were shared with Psm3. In contrast, Psm3-S-Rad21 was barely detectable along chromosomes with merely 12 peaks identified (Figure 6A). A peak aggregate plot of the peaks in common between Psm3 and Psm3-L-Rad21 illustrates the ability of Psm3-L-Rad21 to associate with chromosomes, but the almost complete absence of Psm3-S-Rad21 at these same locations. These results demonstrate that a long linker allows Psm3-L-Rad21 to load onto chromatin. In contrast the short Psm3-S-Rad21 linker largely prevents *in vivo* cohesin loading.

**Figure 6.**
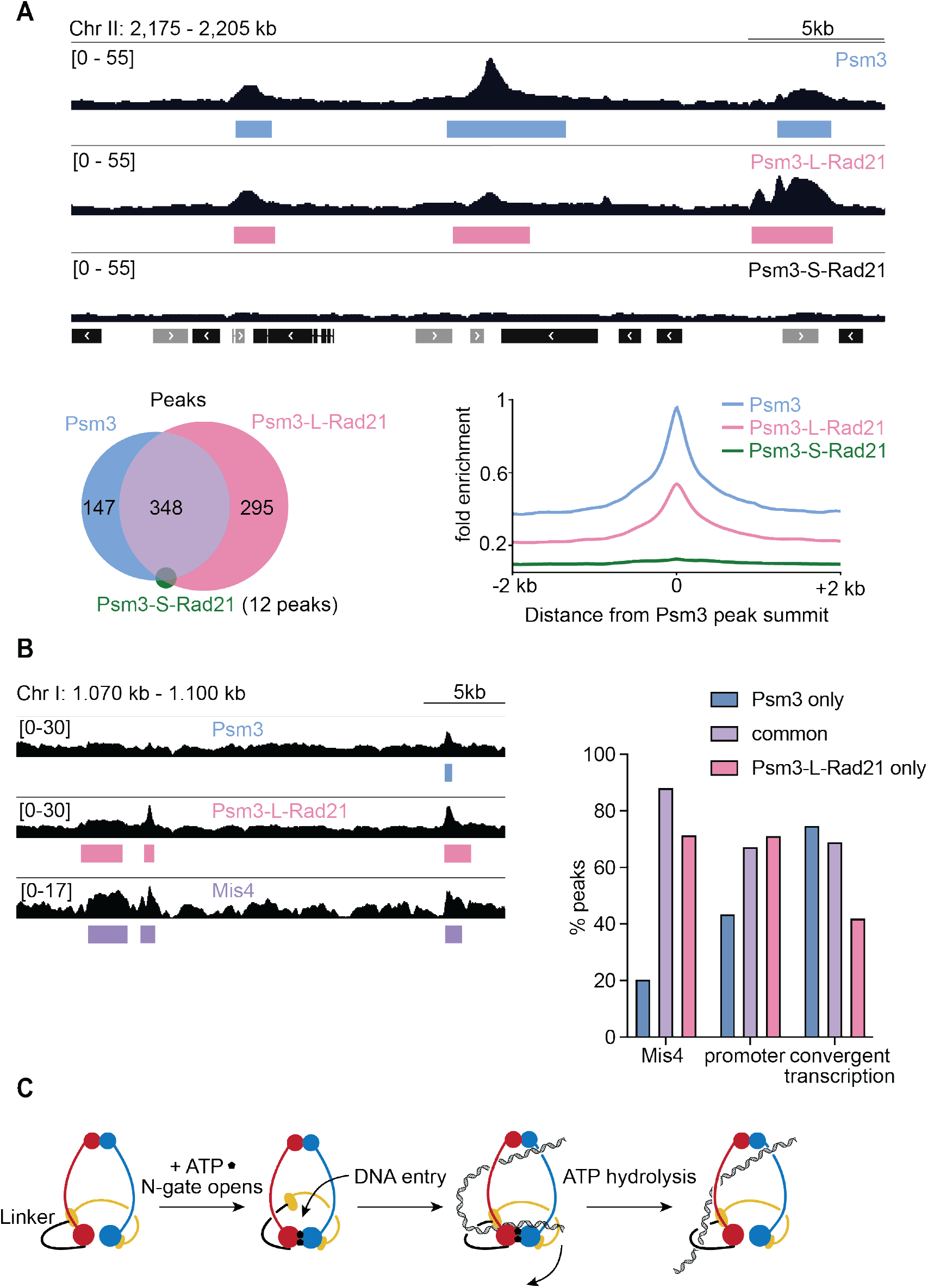
In vivo chromatin loading of N-gate locked cohesin. (A) Calibrated ChIP sequencing traces of Psm3, Psm3-L-Rad21, and Psm3-S-Rad21 along a representative region of chromosome II, with detected peaks demarcated. The positions and orientation of protein coding genes are indicated. The Venn diagram represents detected peak numbers and their overlap, while a peak average plot was generated around the centres of the 348 peaks shared between Psm3 and Psm3-L-Rad21 (Of 12 Psm3-S-Rad21 peaks, 8 were also shared with both, 1 with Psm3-L-Rad21 only, and 3 peaks were unique to Psm3-S-Rad21). (B) ChIP sequencing traces of Psm3, Psm3-L-Rad21, and Mis4 ^43^ along a representative region of chromosome I, with detected peaks demarcated. The percentage of peak positions overlapping with Mis4 binding sites, promoter regions or transcription convergence sites ^33^ is indicated for Psm3 only peaks, Psm3-L-Rad21 only peaks, or those common to Psm3 and Psm3-L-Rad21. (C) Diagram depicting DNA entry into the Psm3-L-Rad21 cohesin ring. The DNA trajectory up to the gripping state (third image from the left) is identical to DNA entry into wild type cohesin. Upon ATP hydrolysis, DNA becomes trapped in the Psm3-L-Rad21 linker compartment, in addition to being entrapped in the cohesin ring.

Most peaks uniquely enriched for Psm3 were at sites of convergent transcription. In contrast, the 295 new Psm3-L-Rad21 peaks that were not shared with Psm3 often coincided with binding sites of the Mis4-Ssl3 cohesin loader ^33,43^ at gene promoter regions. Peaks that were in common between Psm3 and Psm3-L-Rad21 were distributed between cohesin loader binding sites, promoter regions and sites of convergent transcription (Figure 6B). The Psm3-L-Rad21 association pattern could arise for two reasons. Psm3-L-Rad21 cohesin might be recruited by the cohesin loader to promoter regions but then be only slowly or inefficiently loaded. Alternatively, Psm3-L-Rad21 could be loaded by Mis4-Ssl3 at promoter regions but then be impeded from moving away from its loading sites by RNA polymerase II transcription ^31,34^. Future experiments will be required to distinguish between these possibilities. We note that C-gate locked cohesin showed a similar tendency to accumulate at cohesin loader binding sites and promoters (Figure S2B).

The observation that a short Psm3-S-Rad21 linker blocks almost all detectable cohesin loading onto chromosomes suggests that *in vivo* DNA entry into the cohesin ring occurs through the N-gate that is blocked by this linker. If DNA enters cohesin through the N-gate, how can Psm3-L-Rad21 cohesin complexes function, in which the N-gate entry route is also covalently obstructed? As illustrated in Figure 4B, Psm3-Rad21 linkers do not in fact block opening and closing of the N-gate. Furthermore, if the linker peptide is long enough, DNA can be accommodated in a new cohesin compartment that is created by the linker peptide. Consequently, DNA will be trapped both inside the cohesin ring, as well as inside the linker loop (Figure 6C). The linker presence might reduce the cohesin loading efficiency, as observed, and/or impede transcription-driven translocation following loading. Either possibility could explain the additionally detected Psm3-L-Rad21 peaks.

### N-gate, but not hinge gate, operation is linked to the cohesin ATPase

To investigate how the cohesin ATPase operates DNA entry gates, we returned to *in vitro* observations of cohesin behaviour. Structural and biophysical studies showed how the cohesin N-gate opens upon ATPase head engagement, and closes again upon DNA and cohesin loader arrival, providing a molecular framework for this DNA entry route into the cohesin ring ^15,20^. If the cohesin hinge serves as an alternative, or additional, DNA entry gate, one would expect that the hinge gate also responds to the ATP binding state of the cohesin ATPase. To investigate this possibility, we employed fluorescence resonance energy transfer (FRET) to assess the conformational dynamics of all three possible cohesin DNA entry gates.

We inserted SNAP and CLIP tags at cohesin’s three ring subunit interfaces, which we labelled with Dy547 and Alexa 647 dyes respectively (Figure 7). The fluorophore pairs of these double labelled cohesin complexes all displayed measurable proximity, based on the recorded FRET signal between the donor and acceptor dyes. We then incubated the three cohesin gate reporters with or without added ATP, DNA, or cohesin loader and measured resultant FRET changes. As previously observed ^15^, ATP addition in isolation, or together with DNA or the cohesin loader, resulted in FRET reduction at the N-gate, suggestive of gate opening. Addition of ATP together with both DNA and the cohesin loader did not result in a similar FRET reduction, conditions under which the cohesin loading reaction comes to completion and the N-gate ends up closed. In contrast to our observations at the N-gate, ATP addition only marginally affected C-gate FRET. No changes were observed when adding ATP together with any other components. Finally, we investigated the conformational dynamics of the hinge gate. The hinge gate fluorophore pair created a strong FRET signal, but its intensity remained unresponsive to ATP or any combination of added DNA entry reaction components. These results suggest that, of cohesin’s three gates, the N-gate is most responsive to ATP-dependent conformational changes. As cohesin loading onto DNA is an ATP-driven molecular event ^37,38^, the N-gate emerges as the most likely ATP-operated DNA entry route into the cohesin ring.

**Figure 7.**
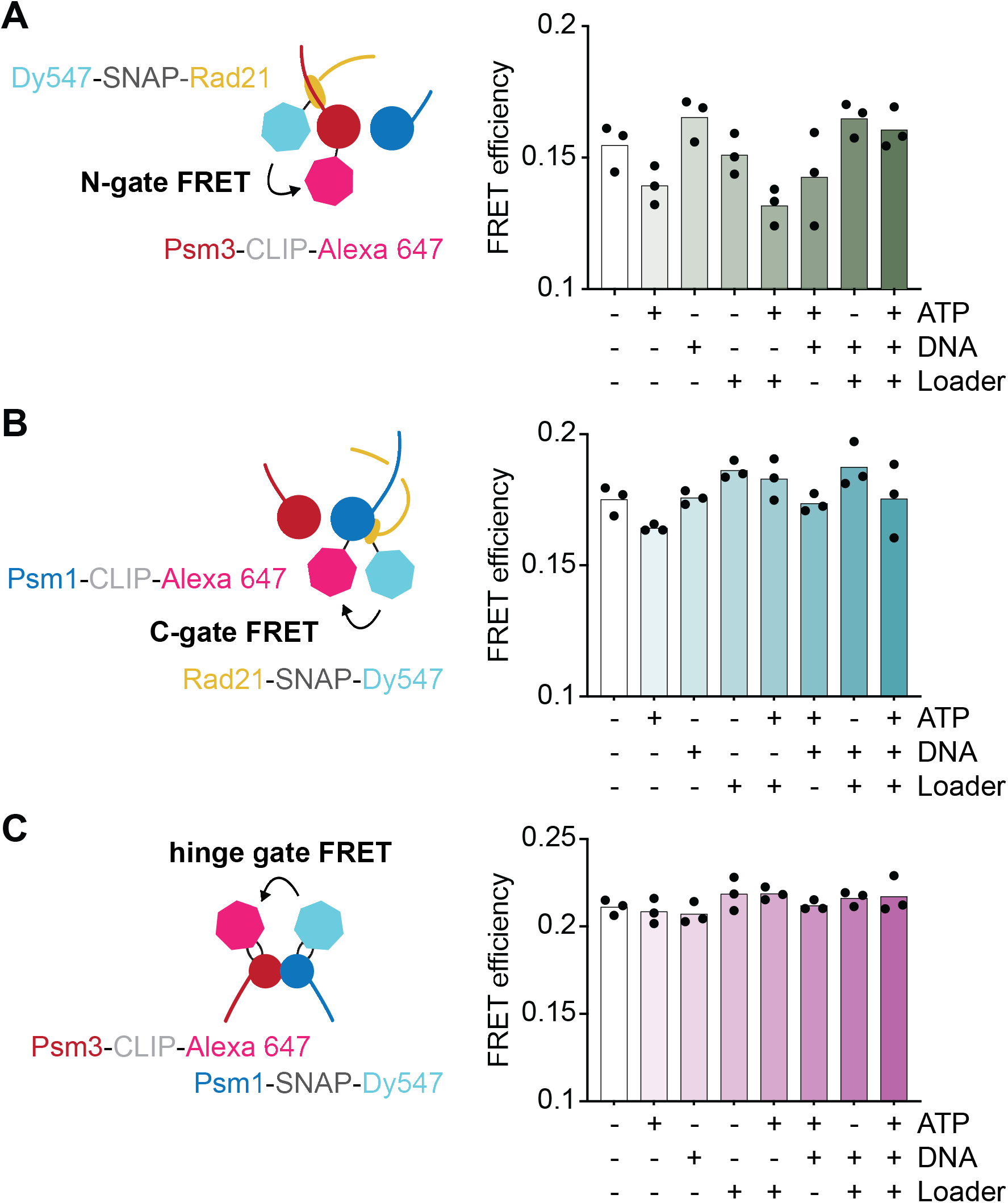
N-gate operation is coupled to the cohesin ATPase. (A) Fluorophore labelling schematic of cohesin’s N-gate. FRET efficiency I_A_/(I_D_ + I_A_) was recorded in presence of the indicated added components, where I_D_ is the donor and I_A_ the acceptor emission intensity resulting from donor excitation. Individual results from 3 independent experiments are shown, the bars represent the means. (B) as (A), but FRET was recorded at the C-gate. (C) as (A), bit FRET was recorded at the hinge gate.

## DISCUSSION

In this study, we investigated DNA entry into the fission yeast cohesin ring. By systematically locking three potential DNA entry gates, we assessed their respective contributions to *in vitro* DNA entry. We also provided an analysis of how the N-gate contributes to *in vivo* cohesin loading. Our approaches reveal that fission yeast cohesin can entrap DNA through more than one of its gates *in vitro*, yet *in vivo*, cohesin’s N-gate emerges as the primary DNA entry route into the cohesin ring.

### *In vitro* DNA entry through more than one gate

Studies with budding yeast cohesin have identified both the N-gate, as well as the hinge gate, as possible *in vitro* DNA entry routes ^19^. Our results suggest that fission yeast cohesin behaves in a similar way, it topologically entrapped DNA if either the hinge gate or the N-gate were locked. Covalent hinge gate closure did not compromise *in vitro* loading onto DNA, while N-gate closure reduced the loading efficiency by around half. The consequence of N-gate closure became more pronounced *in vivo*. Closing the gate with a long peptide linker reduced loading, while a shorter linker abolished cohesin loading onto chromatin.

If, as these results suggest, the N-gate is cohesin’s primary *in vivo* DNA entry route, how can DNA enter the cohesin ring *in vitro* when the N-gate is closed? The hinge is the cohesin ring’s weakest interface ^42^, spontaneous hinge opening has been observed during high-speed atomic force microscopy experiments ^44^, and its measured *in vitro* dissociation half-time is around 15-30 minutes ^12^. If cohesin engages with DNA, but topological DNA entry through the N-gate is prevented by a peptide linker, spontaneous hinge opening could let the DNA in. The duration of *in vitro* DNA loading incubations exceed the hinge dissociation half-time ^8,19^. In contrast, less time is available *in vivo* between cohesin loading in late G1 and sister chromatid cohesion establishment during S-phase ^45^. A chromatin environment might additionally obstruct DNA entry through the hinge.

Cells are inviable if the N-gate is blocked by a short peptide linker but remain viable if a longer linker is used. As we have seen above (Figures 4B, 6C), a long linker does not prevent operation of the N-gate and, while the loading efficiency is reduced, DNA likely reaches a state in which it is embraced by both the cohesin ring and the linker peptide.

### The N-gate as conserved DNA entry gate into SMC complexes

N-gate opening in response to ATPase head engagement is a conserved feature amongst SMC complexes, not only cohesin ^15,20^, but also condensin ^46^, the Smc5-Smc6 complex where the Nse5-Nse6 loader assists N-gate opening ^47^, as well as the bacterial MukBEF complex ^48–50^. Upon DNA arrival, in all cases, the N-gate shuts again. ATP and DNA bound loading intermediates, equivalent to cohesin’s ATP-bound DNA gripping state, in which DNA finds itself locked between the N-gate and the ATPase heads ^15^, have been observed with all studied SMC complexes ^47,49–53^. From there, ATP hydrolysis will lead to head disengagement, allowing DNA to complete a shared trajectory through the N-gate into the SMC ring.

It has been argued that DNA does not in fact topologically enter the condensin ring ^52^. However, this conclusion stems from experiments that were performed with condensin whose N-gate was locked with a peptide linker. Future experiments will have to settle whether DNA topologically enters the condensin ring if the N-gate is left unobstructed. A peptide linker that closes the N-gate was also employed to study the Smc5-Smc6 complex ^47^. In this case, DNA demonstrably became trapped inside the linker compartment following entry through the N-gate. Taken together, available evidence points to the N-gate as the conserved DNA entry path into SMC rings. To date, no molecular mechanism has been proposed by which the SMC ATPase heads would alternatively operate the hinge gate.

We note that the N-gate is not only the entry gate into, but also the DNA exit gate out of the cohesin ring ^23,25^. Entry and exit reactions through the same gate are controlled by the Mis4-Ssl3 loading and Pds5-Wapl unloading factors ^54^. How these factors give directionality to DNA passage through the N-gate remains to be fully understood.

### A role of the cohesin hinge in sister chromatid cohesion establishment?

If ATP-dependent entry into the cohesin ring takes DNA through the N-gate, this trajectory does not exclude a function for the hinge gate at a later stage. Structural studies have found hinges from different SMC complexes in slightly ajar positions ^16,55^, and various SMC hinges share positively charged central channels, though none of them wide enough to accommodate double stranded DNA ^12^. Instead, SMC hinges show affinity for single stranded DNA ^16,56^, and a recent study suggests that SMC hinges embrace, and can even let pass, single stranded DNA ^57^.

The hinge is the cohesin ring’s weakest subunit interface, which breaks at forces in the range exerted by chromosomal translocases, e.g. a replisome ^42^. As the replication machinery approaches cohesin, it can navigate its way through the ring, but more often than not the ring appears to rupture and leave only one of the replication products trapped inside ^58,59^. Might single stranded DNA recognition of the unwound lagging strand template be a mechanism that comes into play in these circumstances? Is single stranded DNA engagement a more widely spread activity of SMC hinges ^60^? While not the subject to ATP-dependent regulation, the role of SMC hinges in DNA transactions merits further investigation.

## Acknowledgements

We thank the Crick Genomics and Fermentation Science Technology Platforms for their support, the Nurse lab for reagents, and all our laboratory members for discussions and comments on the manuscript. This work was supported by a Wellcome Trust Investigator Award (220244/Z/20/Z) and the Francis Crick Institute that receives its core funding from Cancer Research UK, the UKRI Medical Research Council, and the Wellcome Trust (cc2137).

## METHODS

### Yeast strains

*S. pombe* and *S. cerevisiae* strains used for the *in vivo* experiments and protein purification are listed in Table S1. *S. pombe* strains were constructed by either lithium acetate transformation or mating followed by tetrad dissection or random spore selection. *S. cerevisiae* strains were of w303 background and PCR-based gene targeting or plasmid integration were performed following lithium acetate transformation. Colonies were selected based on the presence of auxotrophic or antibiotic resistance markers. Colonies were further tested, as applicable, by polymerase chain reaction-based genotyping, Sanger or Nanopore sequencing, and Western blotting. Ectopic genes encoding Rad21 or Rad21-Psm1 were inserted into the *rad21-k1* strain background at the *arg1^+^* locus under control of the *rad21* or *adh1* promoters, respectively, to achieve comparable expression levels. Genes encoding for Psm3, Psm3-L-Rad21, or Psm3-S-Rad21 were inserted into the *psm3-602* strain background at *arg1^+^* under control of the *psm3* or *tdh1* promoters, respectively. Psm1 was fused to a 3 x sAID tag for auxin-mediated degradation at its endogenous locus by PCR-mediated gene targeting. The gene encoding OsTIR1F74A was inserted at the *arg3^+^* locus under *adh1* promoter control ^62,63^.

### Cohesin and cohesin loader purification

All protein purifications were performed using an Äkta pure (Cytiva) chromatography system. *S. pombe* WT, C-gate locked, and N-gate locked cohesins were expressed in *S. cerevisiae* cells and purified as previously described by sequential purification steps using IgG-Sepharose or HiTrap NHS-activated HP (Cytiva) conjugated to IgG (Sigma-Aldrich), HiTrap Heparin HP (Cytiva), and Superose 6 Increase 10/300 GL (Cytiva) columns ^8,64^.

Hinge gate locked cohesin was expressed in *S. cerevisiae* and purified as previously described ^15^. On the heparin column, R buffer (20 mM Tris/HCl, pH 7.5, 0.5 mM TCEP, 10% (v/v) glycerol) including 100 mM NaCl and 4 μM SC-Cy5 crosslinker ^15^ was injected and incubated at 25°C for 1 hour. The column was washed clear of the crosslinker, cohesin was eluted, and incubated overnight at 4°C to allow for completion of SNAP-CLIP coupling. Peak fractions were concentrated by centrifugal ultrafiltration and applied to a Superose 6 Increase 10/300 GL column. Cohesin was eluted in R buffer containing 200 mM NaCl.

Cohesin used for bulk FRET experiments was expressed and purified as previously described ^15^. The peak fractions from the heparin elution in R buffer containing approximately 600 mM NaCl were concentrated by centrifugal ultrafiltration, supplemented with 2 μM BG-surface Alexa 647, 1 mM DTT, and 0.003% Tween20, and incubated at 25°C for 1 hour. Next, cohesin was supplemented with 4 μM BC-surface Dy547 and incubated at 4°C for 16 hours. The labelled cohesin was applied to a Superose 6 Increase 10/300 GL column and eluted in R buffer containing 200 mM NaCl and 0.003% Tween20.

The *S. pombe* Mis4-Ssl3 cohesin loader was overexpressed in *S. pombe* and purified as previously described by sequential steps on HiTrap NHS-activated HP column (Cytiva) conjugated to IgG (Sigma-Aldrich), HiTrap Heparin HP (Cytiva), and Superdex 200 10/300 GL (Cytiva) columns ^8,64^.

### Cohesin loading assay

Cohesin loading assays were performed as previously described with minor modifications ^8,64^. Cohesin tetramers (100 nM) were mixed with cohesin loader (100 nM), pBluescript KS II DNA (3.3 nM) or pBluescript KS II DNA linearised by PstI digestion (3.3 nM), and ATP (0.5 mM) on ice in reaction buffer (35 mM Tris-HCl pH 7.5, 15% glycerol, 0.5 mM TCEP, 1 mM MgCl_2_, 45 mM NaCl, 0.003% Tween-20). The final reaction volume was 30 μl. The reaction was incubated for 2 hours at 32°C and stopped by addition of 400 μl cold buffer A (35 mM Tris-HCl pH 7.5, 0.5 mM TCEP, 750 mM NaCl, 0.35% triton X-100). Anti-V5 tag antibody (BioRad) was conjugated to Protein A-coated magnetic beads (Invitrogen), added to terminated reactions, and incubated at 4°C overnight rotating. The beads were washed twice with buffer A, twice with buffer B (35 mM Tris-HCl pH 7.5, 0.5 mM TCEP, 500 mM NaCl, 0.1% Triton X-100), and twice with buffer C (35 mM Tris-HCl pH 7.5, 0.5 mM TCEP, 50 mM NaCl, 0.1% Triton X-100). The sample was then split in 2 equal parts. One part was used to quantify the amount of cohesin retrieved by the beads, and the other to visualise the amount of DNA captured by cohesin. To quantify the amount of captured cohesin, beads were resuspended in 15 μl of sample buffer (100 mM Tris-HCl pH 6.8, 4% SDS, 20% glycerol, 4% β-mercaptoethanol) and heated to 98°C for 5 minutes. Proteins were separated by SDS-PAGE and stained with Quick Coomassie Stain (Neo Biotech). To quantify the amount of DNA captured by cohesin, the beads were resuspended in 12 μl of elution buffer (10 mM Tris-HCl pH 7.5, 50 mM NaCl, 1 mM EDTA, 0.75% SDS, 1 mg/ml proteinase K) and incubated at 50°C for 20 minutes. The recovered DNA was separated by 0.8% agarose gel electrophoresis in TAE buffer and stained with SYBR gold (Invitrogen). Protein and DNA were imaged with ChemiDoc MP Imager (Bio-Rad) and quantified with Fiji software ^65^.

To assess loading of hinge gate locked cohesin onto DNA, a 3 kb linear DNA was prepared and immobilized on magnetic beads as previously described ^15^. To evaluate topological DNA entrapment, DNA was digested using the PstI restriction enzyme.

### Bulk FRET measurements

All bulk FRET measurements were carried out at room temperature in reaction buffer (35 mM Tris-HCl pH 7.5, 0.5 mM TCEP, 25 mM NaCl, 1 mM MgCl_2_, 15% (w/v) glycerol and 0.003% (w/v) Tween 20). 40 μl of reaction containing 10 nM Dy547 and Alexa 647-labeled cohesin, 100 nM Mis4-Ssl3, and 3 nM DNA were mixed, and the reaction was started by addition of indicated supplemental components. ATP was used at a final concentration of 0.5 mM. The reactions were incubated at 32°C for 20 minutes. The samples were applied to a 384-well plate and fluorescence spectra were collected on a CLARIOstar high performance plate reader. Samples were excited at 525 nm and emitted light was recorded between 560 - 700 nm in 0.5 nm increments. To evaluate FRET changes caused by cohesin’s conformational changes across different experimental conditions, we report relative FRET efficiency, I_A_/(I_D_ + I_A_), where I_D_ is the donor emission signal intensity at 565 nm resulting from donor excitation at 525 nm and I_A_ is the acceptor emission signal intensity at 665 nm resulting from donor excitation at 525 nm.

### Structural models

Protein structure predictions were generated using the AlphaFold Server, which uses the AlphaFold 3 model ^61^. Figures were created using PyMOL Molecular Graphics System, Version 3.1.6.1 Schrödinger, LLC. Psm3 was coloured in firebrick, Psm1 in marine, Rad21 in yellow orange, Mis4 in grey50, and protein linkers in black.

### Western blot

Around 10^8^ cells were collected, resuspended in 1 ml cold 20% TCA, and kept on ice for at least 10 minutes. Cells were washed with 1 ml 1 M Tris base and resuspended in 100 μl sample buffer (100 mM Tris-HCl pH 6.8, 4% SDS, 20% glycerol, 4% β-mercaptoethanol). Next, cells were heated to 95°C for 5 minutes and then broken at 4°C using a FastPrep-24 bead beater in 4 x 15 seconds cycles at 5 m/s with 5 minutes pauses. Supernatant was collected and the amount of protein quantified using Bradford reagents. 20 μg of protein was separated by SDS-polyacrylamide gel electrophoresis. Proteins were transferred to nitrocellulose membranes, incubated with primary antibody, followed by a secondary HRP-conjugated antibody, both diluted in 5% dried milk in PBSA with 0.1% Tween 20. The antibodies used are listed in the key resources table. Membranes were developed using ECL prime reagents (GE Healthcare) and visualised using a ChemiDoc MP imager (Bio-Rad).

### Viability assay

Cells were plated on YE5S plates in 10-fold serial dilutions. Temperature sensitive strains were incubated at 25°C for 3 days or at 34°C for 2 days. Other strains were incubated at 30°C for 2 days. Strains containing sAID tag and OsTIR1F74A were incubated for 3 days at 30°C on YE5S or YE5S supplemented with 100 nM 5-Adamantyl-IAA (Tokyo Chemical Industry). The plates were imaged on the ChemiDoc MP imager (Bio-Rad) using its colorimetric detection setting.

### Sister chromatid cohesion assay

*S. pombe* strains used to assess sister chromatid cohesion contain tandem repeats of LacO sequences proximal to the centromere of chromosome II and express a LacI-GFP fusion protein, which binds to LacOs ^28,29^. Temperature sensitive strains were grown in YE5S media overnight at 25°C and transferred to 37 °C for 6 hours. AID strains were grown in YE5S media overnight at 30°C, 100 nM 5-Adamantyl-IAA (Tokyo Chemical Industry) was added and samples taken after 1 and 6 hours. 2 ml of culture was collected. Cells were harvested by centrifugation and fixed in 100% ethanol overnight at-20°C. Cells were washed in PBSA and mounted on a 2% agarose patch on a microscope slide. Cells were imaged and GFP signals were detected using a DeltaVision Olympus IX70 inverted microscope equipped with a 100x (NA = 1.40) PlanApo objective. z-stacks with 35 images at 0.2 μm intervals were acquired, deconvolved, and merged using maximum intensity projection. 100 cells were analysed for each sample.

### Quantitative ChIP-sequencing

*S. pombe* cells were grown in YE5S overnight at 25°C. Per sample, 10^9^ cells were crosslinked by addition of 1% formaldehyde for 30 minutes, then the crosslinking reaction was quenched with 150 mM glycine for 10 minutes. Cells were harvested by centrifugation and washed with cold PBS. *S. cerevisiae* cells, used for spike-in normalisation were grown in YPD overnight at 30° C. Per sample, 2×10^8^ cells were crosslinked by addition of 1% formaldehyde for 20 minutes and the crosslinking reaction was quenched with 150 mM glycine for 10 minutes. Cells were harvested by centrifugation and washed twice with cold PBS and resuspended in 800 μl lysis buffer (50 mM HEPES-KOH pH 7.5, 1 mM EDTA pH 8, 150 mM NaCl, 1% Triton X-100, 0.1% sodium deoxycholate, 1 mM PMSF, 2 x cOmplete EDTA-free protease inhibitor cocktail (Roche)). *S. pombe* cells were broken at 4°C in a FastPrep-24 bead beater using twelve 20 second cycles at 5.5 m/s with 5 minutes pauses. *S. cerevisiae* cells were broken at 4°C in a Yasui Kikai Multi-Beads Shocker using twenty-eight 60 seconds ON 60 seconds OFF cycles at 2,500 rpm. Cell lysates were collected, beads washed with 400 μl lysis buffer, and the lysates combined. Lysates were centrifuged for 10 minutes at 13,300 rpm at 4°C, the supernatant was discarded, and the pellet was washed with 800 μl lysis buffer. Now *S. pombe* and *S. cerevisiae* samples were mixed at a cell equivalent ratio of 5: 1. The samples were centrifuged for 10 minutes at 13,300 rpm at 4°C, supernatants discarded, and pellets resuspended in 750 μl lysis buffer without PMSF or protease inhibitors. Chromatin was sheared by sonication to an average size of 300 - 500 bp using 40 cycles of 30 seconds ON 30 seconds OFF in a Bioruptor Plus (Diagenode) at its high setting. Sonicated samples were centrifuged, the supernatants collected, and protein concentrations quantified using Bradford reagents. 50 μl input chromatin sample was collected. Chromatin samples equivalent to 5 mg of protein were used for chromatin immunoprecipitation using anti-V5 tag antibody coated Protein A magnetic beads at 4°C overnight. Beads were washed two times in lysis buffer without PMSF or protease inhibitors, two times in lysis 500 buffer (50 mM HEPES-KOH pH 7.5, 1 mM EDTA pH 8, 500 mM NaCl, 1% Triton X-100, 0.1% Na-Doc), two times in LiCl buffer (10 mM Tris-HCl pH 8, 1 mM EDTA pH 8, 250 mM LiCl, 1% IGEPAL, 1% sodium deoxycholate), and one time in TE (10 mM Tris-HCl pH 8, 1 mM EDTA). Beads were resuspended in 200 μl elution buffer (50 mM Tris-HCl pH 8, 10 mM EDTA, 1% SDS). Input samples were resuspended in 150 μl input elution buffer (10 mM Tris-HCl pH 8, 1 mM EDTA, 1% SDS). Reverse crosslinking was performed at 65°C overnight on a thermomixer shaking at 1,200 rpm of both chromatin immunoprecipitate and input samples. Next, DNA was extracted by phenol-chloroform extraction, and the aqueous fraction was further purified using the DNA Clean & Concentrator-5 kit (Zymo Research). Sequencing libraries were constructed with the NEBNext Ultra II DNA Library Prep Kit for Illumina (New England Biolabs). Paired-end sequencing of ∼10 million reads per sample was performed on a NovaSeq X Series sequencer (Illumina).

### ChIP-sequencing data analysis

Sequencing data was processed using a custom fork of the nf-core/chipseq pipeline to handle spike-ins ^66^. The pipeline was executed using a custom Singularity profile ^67^. Raw reads were assessed with FastQC. Adapter sequences, primers, poly-A tails, and low-quality bases were removed using Trim Galore. Trimmed reads were aligned to a hybrid genome made of *S. pombe* genome ASM294v2 ^68^ and *S. cerevisiae* genome sacCer3 ^69^ using BWA ^70^. Aligned reads were processed with Picard to mark PCR duplicates and produce BAM files. Spike-in scale factors were calculated for each sample from the fraction of reads mapping to the *S. cerevisiae* genome: scale_factor = 1 × 10⁶ / spikein_reads. Genome-wide coverage bigWig files were generated with BEDTools ^71^ (bedGraphToBigWig), normalised to 1,000,000 mapped reads (RPKM), and scaled by the spike-in scale factor. Genome-wide IP enrichment relative to matched input controls was calculated with deepTools ^72^ (bamCompare). Normalised strand cross-correlation (NSC) and relative strand cross-correlation (RSC) quality metrics were performed with phantompeakqualtools.

Peaks were called in all samples with MACS3 ^73^ tool run in broad peak mode with --macs_fdr = 0.000001 and --broad_cutoff = 0.0001. Consensus peak set across replicates was generated with BEDTools ^71^ by merging overlapping peak coordinates. Peaks were called only if they were present in at least 2 replicates. Peaks longer than 10 kb and located at centromeres were manually removed. Reads within consensus peaks were counted with featureCounts ^74^. Read counts across consensus peaks were quantified using DiffBind ^75^ (R, Bioconductor) by counting reads within 500 bp windows centred on peak summits. Principal component analysis (PCA) was performed on the normalised read count matrix.

Peaks were classified using a custom BED annotation file that contained tRNA and promoter (±500 bp window around the transcription start site of each protein-coding gene) features derived from the PomBase GFF3 genomic sequence features ^76^, supplemented by convergent transcription sites and Mis4 peaks ^35^. A peak was considered to overlap an annotated region if a minimum of 1 bp of the peak interval intersected the feature. If a peak overlapped with multiple features, a feature whose boundary was the closest to the peak’s summit was assigned to the peak. Convergent transcription sites were identified from the PomBase GFF3 genomic sequence features ^76^. All genes were sorted by chromosomal start position and scanned pairwise to identify adjacent gene pairs in a convergent orientation. To obtain Mis4 peaks, we reanalysed 6 samples of calibrated ChIP-seq of *S. cerevisiae* Scc2, where *S. pombe* cells containing Mis4-Myc were added as a spike-in control ^35^. Sequencing data was processed as described above. Spike-in normalisation was not applied.

Genome-wide signal tracks for visualisation and cross-condition comparison were generated using both IP and matched input control BAM files with MACS2 (v2.2.9) ^73^ in merged mode – all replicates BAM files for each condition were pooled into a single peak-calling run in broad peak mode generating signal per million reads (SPMR) normalised bedGraph output. SPMR-normalised bedGraph files were converted to bigWig format using bedGraphToBigWig (UCSC Kent tools) ^77^. The SPMR bigWig files were further normalised using the spike-in scale factors. Per-replicate scale factors were combined into a pooled scale factor for the merged sample:

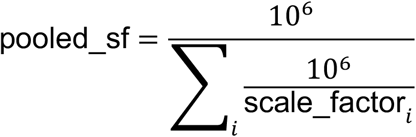

The SPMR bedGraph for each sample was multiplied by its pooled scale factor and the result was converted back to bigWig.

Peak average plots were generated using deepTools ^72^ (computeMatrix and plotProfile). For each comparison between conditions, peaks common to both conditions were identified by reciprocal 1 bp overlap. Within each common peak, the position of maximum signal in the reference condition’s (Psm3 or Rad21) bigWig was used as the peak summit. Coverage matrices were computed in reference-point mode, centred on peak summits with a ±2 kb flanking window. Average signal profiles were plotted for each condition using plotProfile.

---

**Figure S1.**
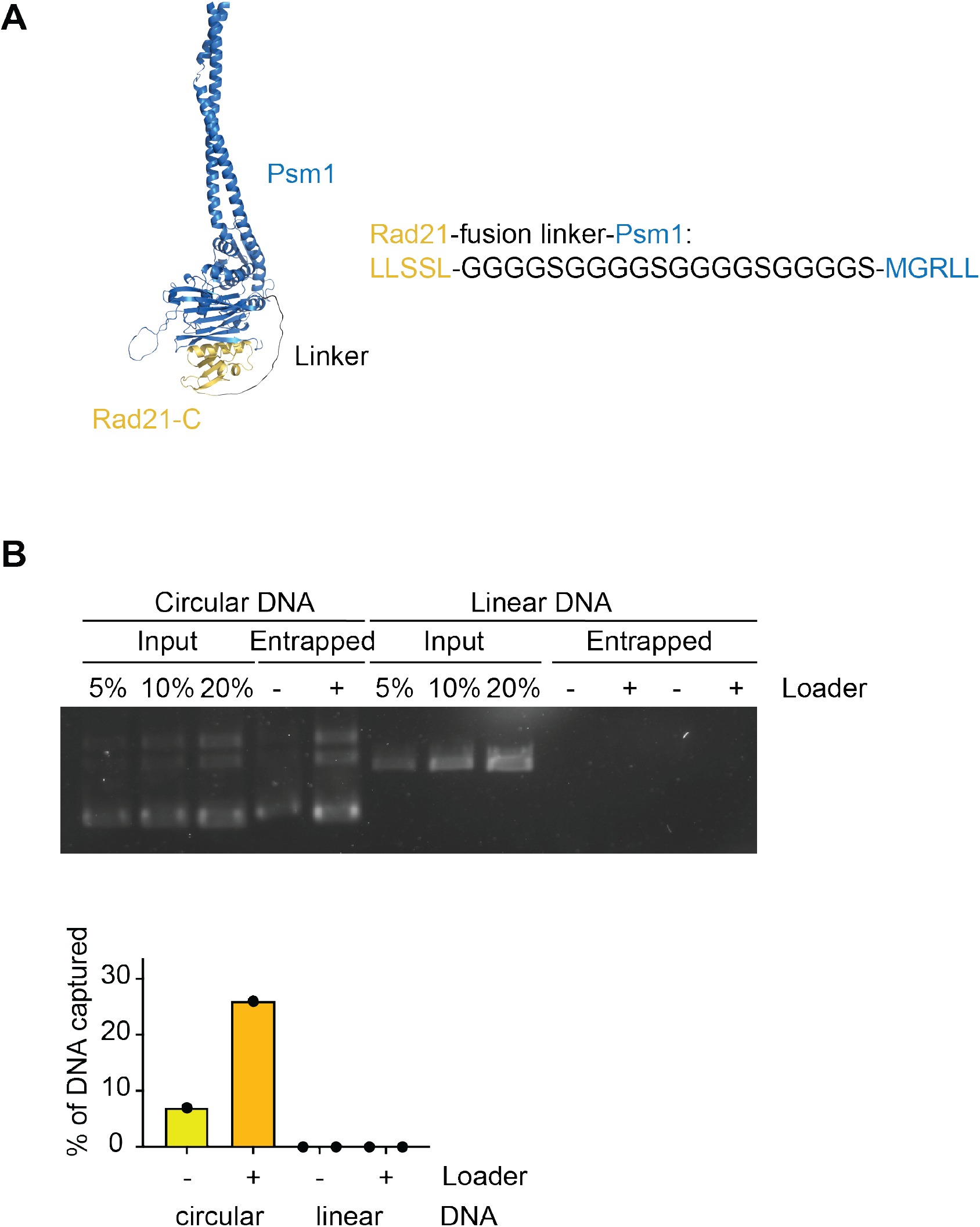
C-gate locked cohesin. (A) Structural model of C-gate locked cohesin. The linker peptide sequence used to fuse the Rad21 C-terminus to the Psm1 N-terminus is shown. Structural model generated using AlphaFold 3 ^61^. (B) Agarose gel image of DNA recovered in a WT cohesin loading assay using circular or linear DNA, with and without added cohesin loader. Recovered DNA was quantified and the individual results are shown. Two technical repeats of the experiment with linear DNA were included, to ensure reproducibility.

**Figure S2.**
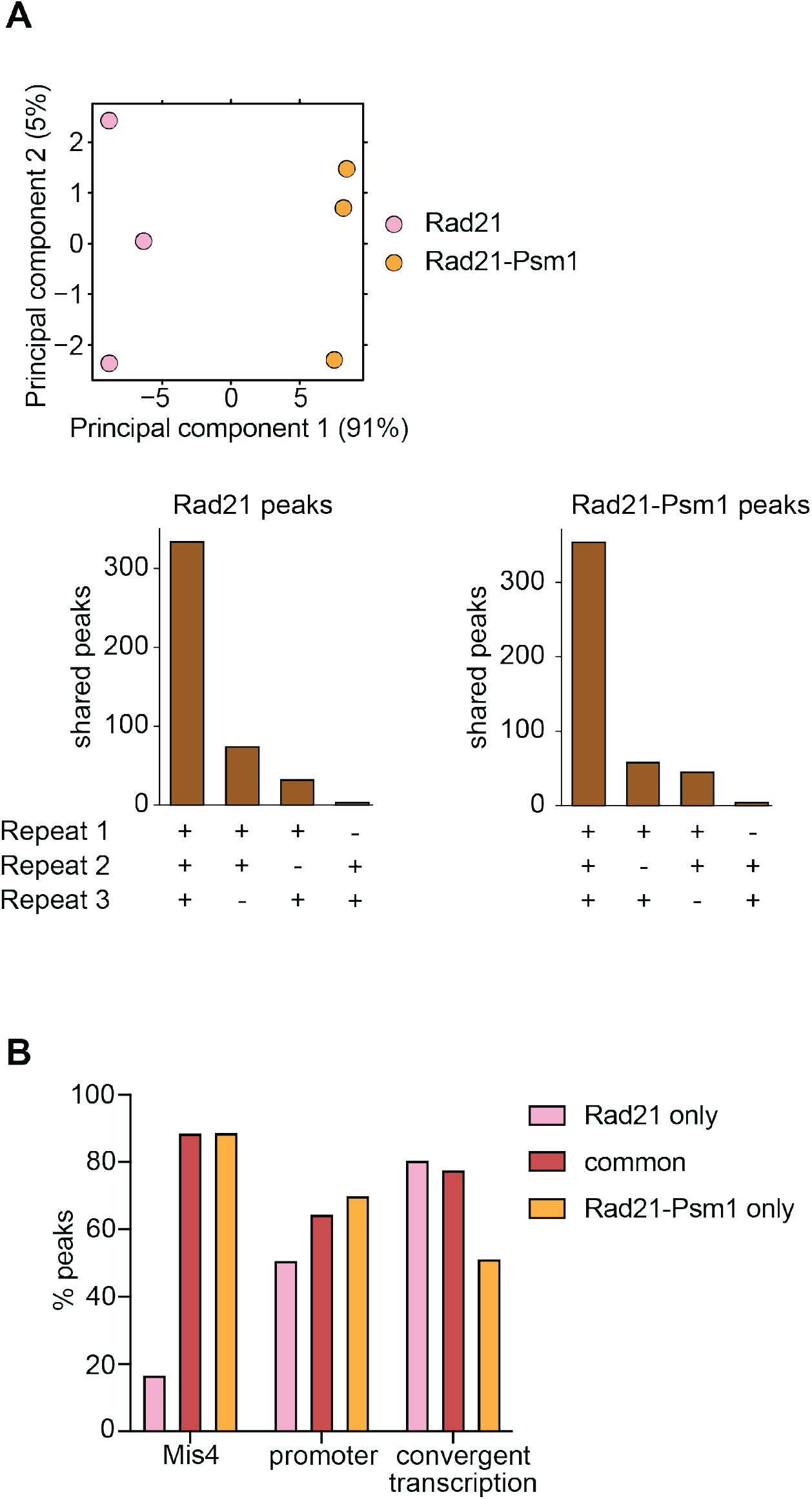
Rad21-Psm1 ChIP sequencing analysis. (A) Principal component analysis of the three independent Rad21 and Rad21-Psm1 ChIP-seq experiments, as well as a breakdown of the Rad21 peak numbers shared amongst the three repeats. (B) Stratification of Rad21 and Rad21-Psm1 peaks by known cohesin binding site features ^33^.

**Figure S3.**
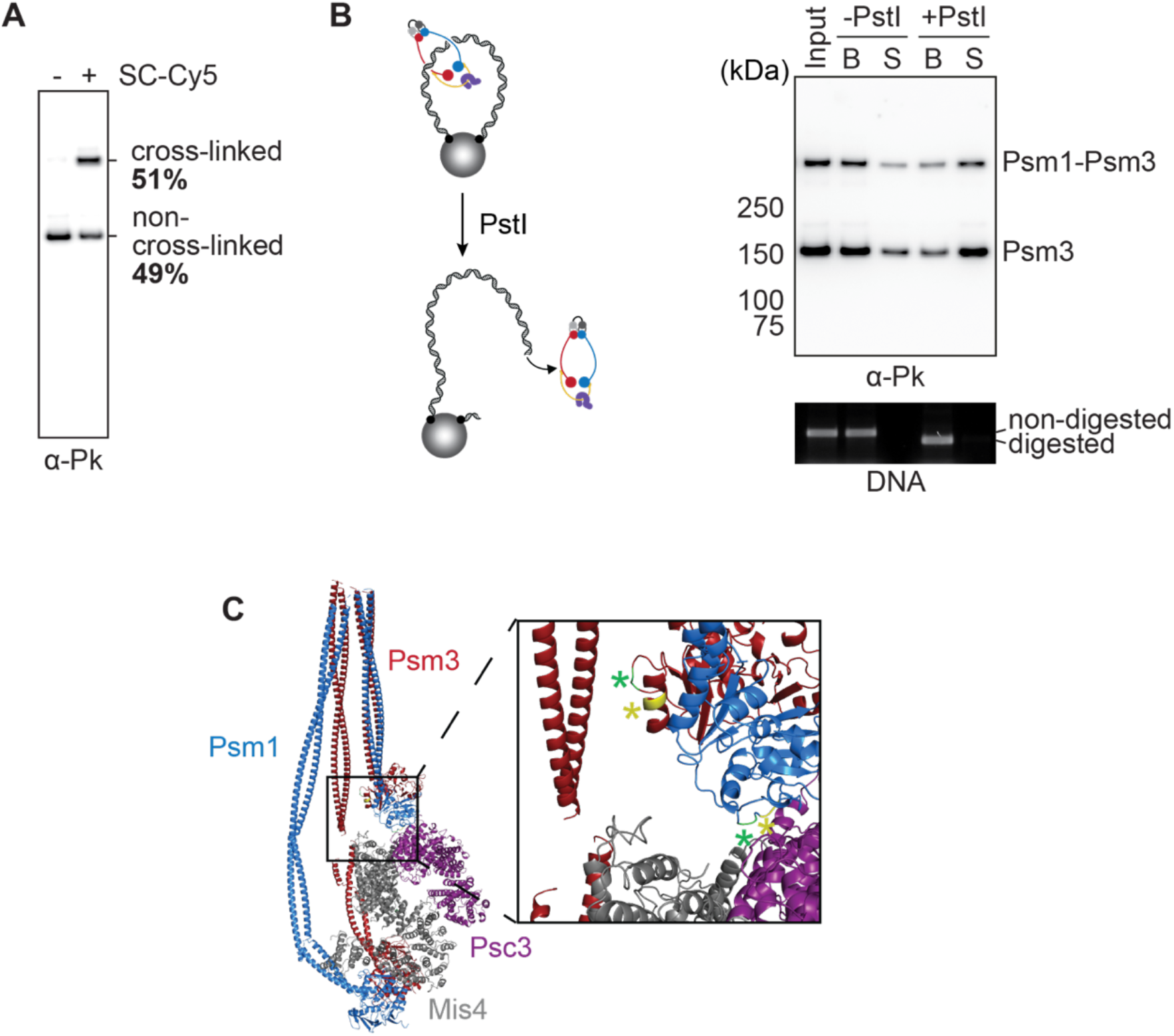
In vitro DNA entry into hinge gate locked cohesin. (A) Cohesin with SNAP-tag and CLIP-tag hinge insertions in Psm1 and Psm3, respectively, was left untreated or was crosslinked with SC-Cy5. Western blot analysis against the Pk-epitope tag on Psm3 was used to quantify the crosslinking efficiency. (B) Schematic of an experiment to test the topological nature of hinge gate locked cohesin loading onto magnetic bead tethered DNA. Following a high salt wash, the DNA was cleaved using the PstI restriction endonuclease and the reaction was split into beads (B) and supernatant (S) fractions. Cohesin was visualised by immunoblotting against the Pk-epitope on Psm3, and the DNA by agarose gel electrophoresis. (C) Structural model of fission yeast cohesin in the DNA gripping state ^40^. The positions of our SNAP and CLIP tag insertions used to lock the hinge gate are shown in yellow, the position used in previous studies is shown in green ^17,19,23^.

**Figure S4.**
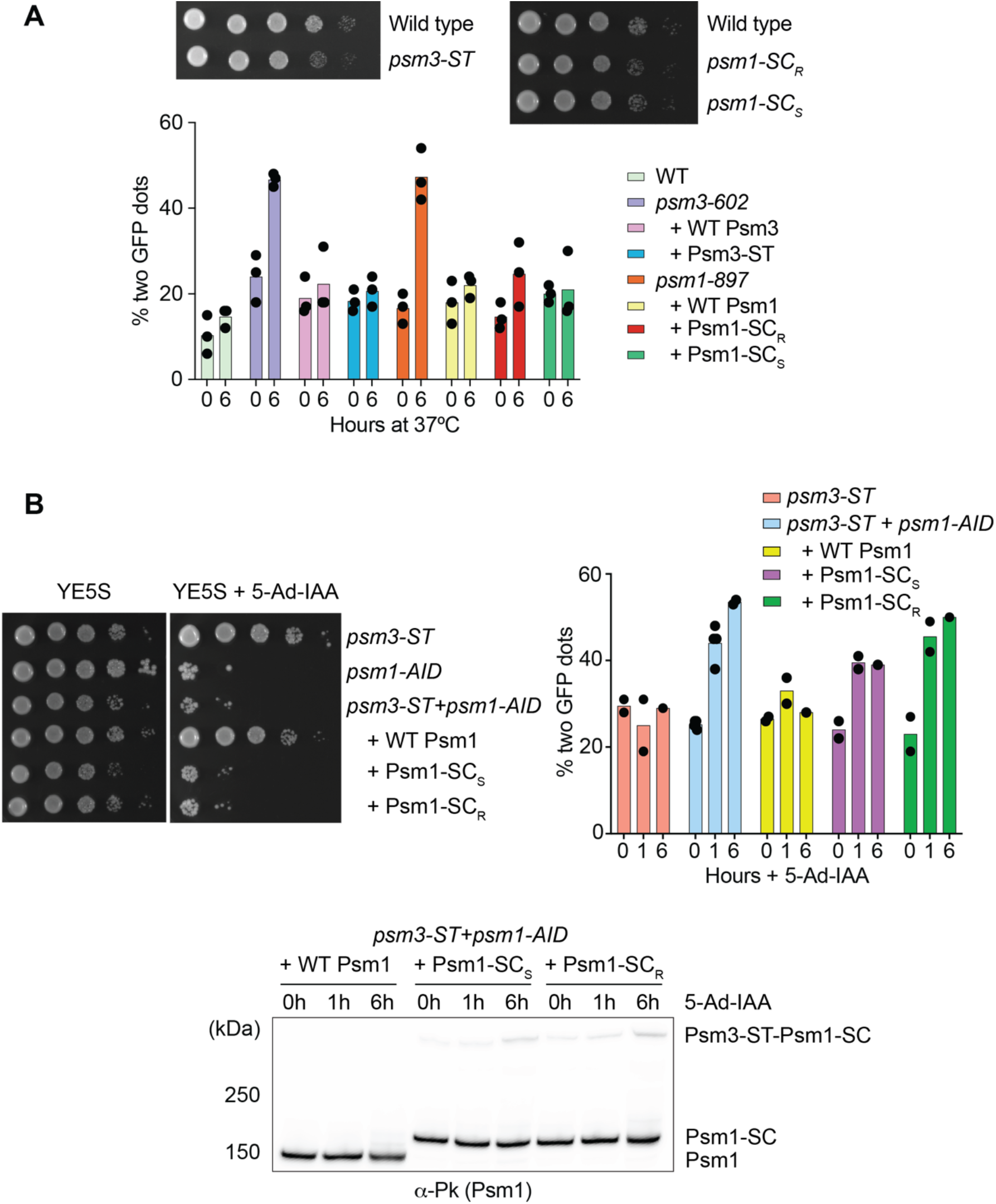
An in vivo approach to locking the cohesin hinge gate. (A) Ten-fold serial dilutions of control cells, cells with a SpyTag inserted at the endogenous *psm3^+^* locus (Psm3-ST, inserted after R585), and cells with a SpyCatcher inserted at the endogenous *psm1^+^* locus (both Psm1-SC_R_ inserted after R640, and Psm1-SC_S_ inserted after S664) were spotted on YE5S plates and grown at 30°C. Quantification of sister chromatid cohesion defects in the same strains is shown, seen as two separated GFP foci, before and 6 hours after shift to 37°C. The individual results of three biological repeat experiments are shown, bars represent the means. (B) The endogenously inserted Psm3-ST was now combined with an ectopic copy of Psm1-SC_R_ or Psm1-SC_S_, in a strain background in which endogenous Psm1 could be depleted via an auxin-inducible degron (*psm1-AID*). Ten-fold serial dilutions of cells with the indicated genotypes were spotted on YE5S plates or YE5S plates containing 100 nM 5-adamantyl-indole-3-acetic acid (5-Ad-IAA) to deplete endogenous Psm1 and grown at 30°C. Quantification of sister chromatid cohesion defects in the same strains is shown, seen as two separated GFP foci, before and 6 hours after 100 nM 5-Ad-IAA addition. The individual results of two biological repeat experiments are shown, bars represent the means. Samples of the culture were taken before, 1 hour and 6 hours after 5-Ad-IAA addition and processed for immunoblotting against the Pk epitope tag (marking Psm1, Psm1-SC_R_ and Psm1-SC_S_) to monitor covalent bond formation between Psm3-ST and Psm1-SC.

**Figure S5.**
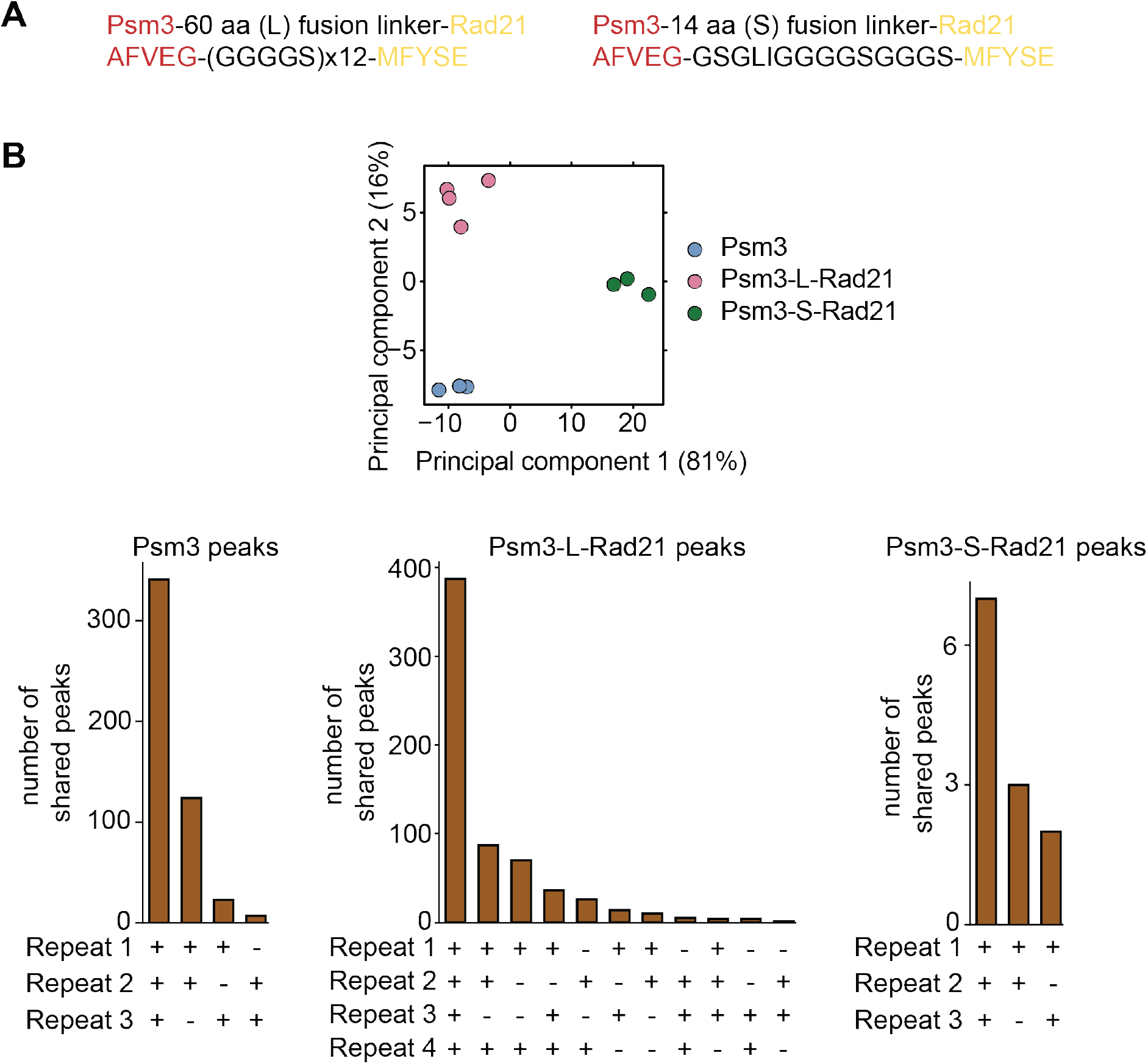
Characterisation of N-gate locked cohesin. (A) Depiction of the Psm3-L-Rad21 and Psm3-S-Rad21 linker sequences. (B) Principal component analysis of at least three independent Psm3, Psm3-L-Rad21, and Psm3-S-Rad21 ChIP-seq experiments, as well as a breakdown of the peak numbers shared amongst the repeats.

**Table S1:**
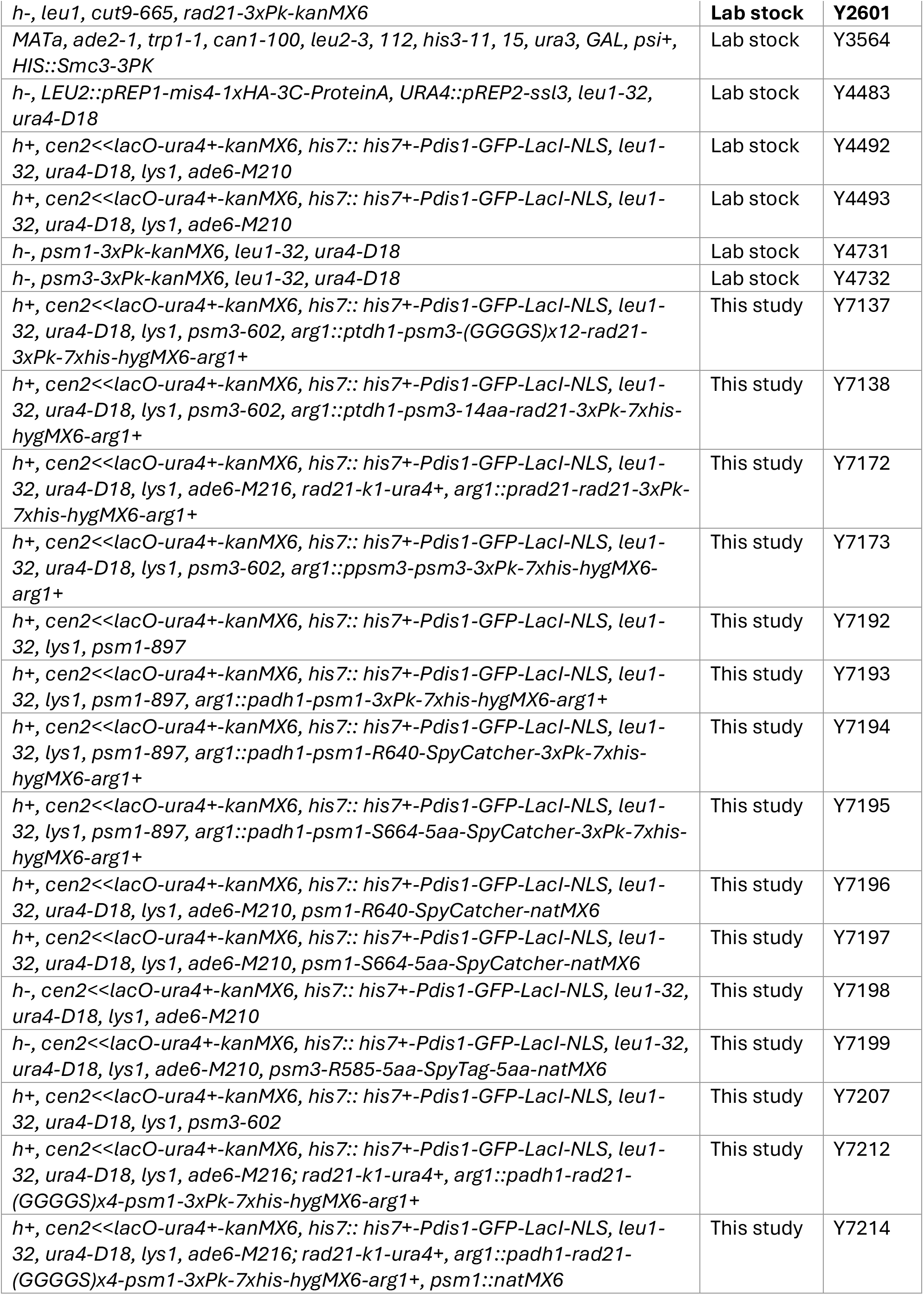
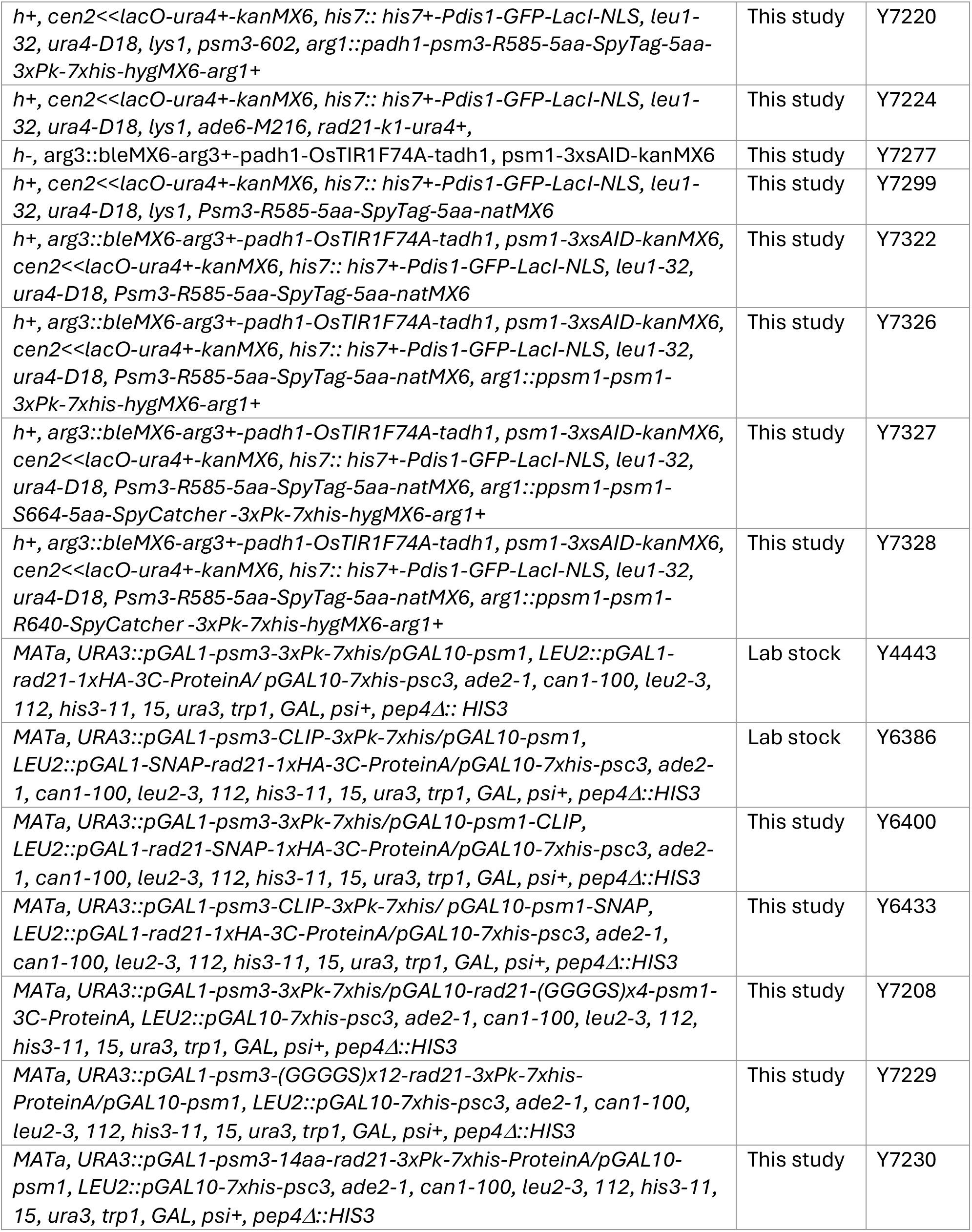
Yeast strains.

